# Mink on the brink: Comparing survey methods for detecting a Critically Endangered carnivore, the European mink *Mustela lutreola*

**DOI:** 10.1101/2022.07.12.499692

**Authors:** Elizabeth Croose, Ruth Hanniffy, Andrew Harrington, Madis Põdra, Asun Gómez, Polly L. Bolton, Jenna V. Lavin, Samuel S. Browett, Javier Pinedo Ruiz, David Lacanal Arnaez, Iñaki Galdos, Jon Ugarte, Aitor Torre, Patrick Wright, Jenny MacPherson, Allan D. McDevitt, Stephen P. Carter, Lauren A. Harrington

## Abstract

Monitoring rare and elusive species is critical in guiding appropriate conservation management measures. Mammalian carnivores are challenging to monitor directly, due to their generally nocturnal and solitary lifestyle, and relatively large home ranges. The European mink *Mustela lutreola* is a Critically Endangered, small, semi-aquatic carnivore and is one of the most threatened mammal species in Europe. In northern Spain, the European mink population is monitored regionally using different methods and approaches, making assessment of national population status difficult. There is an urgent need to 1) assess the efficacy of survey methods, and 2) identify a standard monitoring methodology that can be deployed rapidly and inexpensively over large areas of the mink’s range. We deployed four methods – camera trapping, hair tubes, live trapping, and environmental DNA (eDNA) from water samples – to compare the probability of detecting European mink when present at 25 sampling sites within five 10×10km squares, and the economic cost and time required for each method. All four methods successfully detected European mink but the probability of detection varied by method. Camera trapping and hair tubes had the highest probability of detection; however, eDNA and live trapping detected mink in one 10×10km square where the latter two methods did not. For future European mink monitoring programmes, we recommend a combination of at least two methods, and suggest that camera traps or hair tubes are combined with live trapping or eDNA (depending on the scale and aims of the study), to gather critical information on distribution, occupancy and conservation status.

## INTRODUCTION

Monitoring rare and elusive species is critical in guiding local conservation management measures (Campbell *et al*. 2002, Scheele *et al*. 2019). This is particularly important when monitoring populations or species that are newly reintroduced into the wild, or those that are subject to change, where the reliable and rapid collection of occupancy and distribution data is vital towards establishing both their short- and long-term viability. The success of these monitoring programmes is, however, dependent on the monitoring method used. Key considerations are the behaviour and ecology of the species (Sales *et al*. 2020a) and, since funding will almost always be a limiting factor, the economic cost, effort required, and likely detection success (Barea-Azcon *et al*. 2006, Descalzo *et al*. 2021).

Mammalian carnivores are a particularly challenging group to monitor directly (e.g. by a direct count census) over large spatial scales because they are generally nocturnal, solitary and have relatively large home-ranges (Barea-Azcon *et al*. 2006). The challenge is even greater if the species occurs at low density or is patchily distributed, whether due to natural rarity or population decline. European mink are Critically Endangered, semi-aquatic carnivores, and one of the most threatened mammal species in Europe (Maran *et al*. 2016). European mink were historically widespread across Europe until the 19^th^ century, but their total population size has declined by more than 90% since the beginning of the 20^th^ century (Maran *et al*. 2016) and now probably occupies less than 3% of its former range (Harrington *et al*. 2018). The current distribution of European mink is restricted to small and isolated populations in three areas: Russia; the Danube Delta in Romania and Ukraine; and northern Spain and western France (Maran *et al*. 2016, Zuberogoitia *et al*. 2018). Small populations have also been introduced to Hiiumaa Island, Estonia (Maran *et al*. 2017), Kunashir Island in the Russian Far East (Kisleyko *et al*. 2022) and Lower Saxony, Germany (Lüers & Brandt 2014). The most significant threat to European mink, wherever the species has survived, is competitive exclusion by the invasive American mink *Neovison vison* (Sidorovich 2001, Põdra *et al*. 2013), and, to a lesser extent, habitat loss and non-natural mortality such as road kills (Palazón *et al*. 2012, Zuberogoitia *et al*. 2018, reviewed in Maran *et al*. 2017).

In Spain, the European mink population occupied 1,900-2,000km of watercourses in the north of the country in the early 2000s (Palazón *et al*. 2002). Since then, European mink distribution area has decreased significantly due to the expansion of American mink (Põdra & Gomez 2018) and their total population size is estimated to be around 500 individuals (A. Gómez & M. Põdra, unpublished data 2022). The Spanish population is no longer connected with the smaller French population (estimated at < 250 individuals in 2014; Direction Régionale de l’Environnement *et al*. 2021) and is considered perhaps only the second viable European mink population in its global range, after that in the Danube Delta (Maran *et al*. 2017). American mink are present in one-third of the European mink’s range in Spain, and despite control efforts since the late 1990s and early 2000s (Põdra *et al*. 2013, Mañas *et al*. 2016, Maran *et al*. 2017) their distribution continues to expand (Põdra & Gómez 2018).

Systematic monitoring of European mink is critical for assessing changes in the distribution, abundance and status of their populations. Currently, in Spain, European mink are monitored by regional governments using different methods and approaches, which makes assessing the status of the mink population on a national scale difficult. The presence of American mink presents an extra challenge for monitoring European mink in areas where the two species co-occur as their physical similarities make distinguishing between the two species difficult (footprints detected, or photos/videos taken by camera traps in particular). To date, there has been no scientific comparison of the relative efficacy of the different methods available to monitor European mink and no consensus on which might be the most effective and accurate. As such, there is an urgent need to assess the efficacy of survey methods and provide recommendations on a suitable standard methodology which could be applied across the mink’s Spanish range and elsewhere.

Detection rates vary among methods used to monitor wildlife populations and many methods have not been compared or validated in terms of their efficacy or reliability (Diggins *et al*. 2016, Witmer 2005). Currently, live trapping and camera trapping are the main methods used for monitoring European mink (e.g. Palazón *et al*. 2002, Gómez *et al*. 2011, González-Esteban *et al*. 2004, Põdra 2021), but the relative efficacy of these methods in reliably detecting European mink has never been directly compared. Hair tubes (sections of plastic pipe with adhesive patches to capture hair of animals that enter) have also previously been used to survey European mink and have been widely used for European pine marten *Martes martes*, a similar sized carnivore (Mullin *et al*. 2010). Environmental DNA (eDNA)-based monitoring has not previously been tested for European mink, but has been successful in detecting other carnivores and semi-aquatic species (Sales *et al*. 2020a; Broadhurst *et al*. 2021). The emergence of eDNA has revolutionised biodiversity monitoring in marine and freshwater ecosystems (Deiner *et al*. 2017) and has been successfully applied for detecting a broad range of semi-aquatic and terrestrial mammals in both lentic and lotic systems (e.g. Harper *et al*. 2019, Seeber *et* al. 2019, Sales *et al*. 2020a, b, Broadhurst *et al*. 2021). eDNA metabarcoding has been shown to be comparable to or even outperform traditional survey methods such as field signs or camera traps for monitoring several mammal species (Fediajevaite *et al*. 2021, Sales *et al*. 2020a, b, Lyet *et al*. 2021) and therefore has the potential to be an effective additional method to existing monitoring approaches for European mink (Sales *et al*. 2020a; Broadhurst *et al*. 2021).

The aim of this study was to compare the efficacy of different survey methods for the Critically Endangered European mink with a view to identifying the most suitable candidate method for a standardised survey approach for the species, to provide comparative data to aid in the interpretation of existing monitoring information obtained from various different methods, and, in particular, to explore the potential of eDNA as an additional tool that could be used to provide range-wide distributional data. We deployed four methods – camera trapping, hair tubes, live trapping and eDNA from water samples – in northern Spain on an extant European mink population. We evaluated and compared the probability of detection and the economic cost and time required for each method. Finally, we discuss the advantages and disadvantages of each method and make recommendations for future surveys and monitoring studies.

## MATERIALS AND METHODS

### Study area

The study was carried out in north-eastern Spain in the regions of La Rioja, Álava and Aragon. The European mink has been detected in only a small part of Aragon (Gómez *et al*. 2011) and is less abundant here than in other parts of its range. The status of European mink in La Rioja and Álava has worsened considerably in recent years, due to the invasion of American mink, but successful eradication of American mink during the project LIFE LUTREOLA SPAIN (2014-2019) has enabled European mink to persist there. A small number of captive-born European mink (n=15) were released in La Rioja and Álava in 2018 to reinforce the population in the Ebro basin (http://lifelutreolaspain.com, M. Põdra *et al*., unpub. data).

The surveys took place on the river Ebro, its tributaries and connected wetlands: Salburua wetland, Ea-Tiron, Oja, Najerilla, Leza, Tirón, and Zadorra (Fig. 1). The altitude is between 400-520 m. The Ebro river is approximately 30-100 m wide in this area, with tributaries varying between 5 and 15 m. The area of Salburu wetland is 200 ha. The region is characterised by a continental Mediterranean climate or transition from Mediterranean to Atlantic (Álava) with wetland, lakes, canals and small rivers, poplar groves and narrow riparian forests formed mostly by *Salix sp, Populus sp* and *Alnus glutinosa*, on the river banks. During the study, temperatures at the site ranged from 11 °C to 20 °C. A period of heavy rain occurred between the first and second eDNA water sampling sessions. During this time the Rio Ebro was in flood conditions and water levels rose by over 1 metre.

**Fig. 1.**
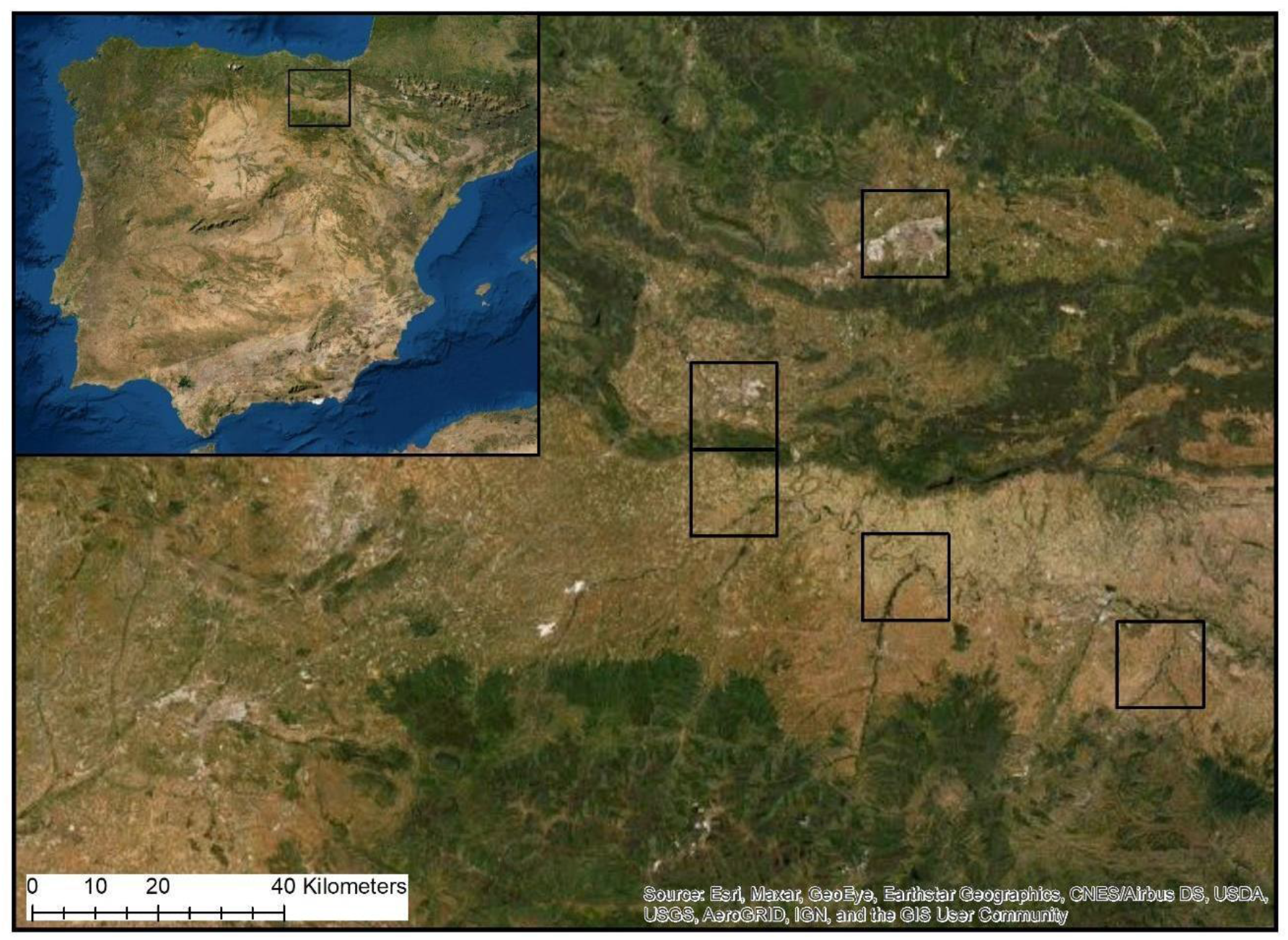
Location of the study area within Spain where four surveying methods were implemented to detect European mink

### Study design

Fieldwork and data collection for all four methods were completed during October-November 2019 and thus overlapped temporally. Data collection for all four methods was carried out in the same five 10 km × 10 km squares based on the Universal Transverse Mercator (UTM) grid reference system. The UTM squares selected represent rivers throughout the whole European mink distribution area, comprising the core area (where European mink are present regularly) and periphery (where European mink are present occasionally). Within each UTM square, five sampling sites were selected, each separated from the adjacent site by approximately 1km of river (spacing selected to maximise the chance of having at least one unit present in a female mink’s home range), giving a total of 25 sampling sites at which the methods were deployed (Fig. 1). Each sampling site was alongside a river or stream bank and sites differed in their characteristics, from main river systems to small streams to marginal backwater.

### Camera trapping

Two camera traps (a Bushnell Trophy Cam HD and a Browning Strike Force HD Pro X) were deployed in a pair at each site. The rationale for deploying two cameras was to maximise the chance of capturing images with a clear view of the mink’s face (necessary for distinguishing between European and American mink) and to compare the efficacy of each camera model. The cameras were deployed either on a stake pushed into the ground, or tied to a tree, at up to 1 m above the ground (Fig. 2a). The cameras were baited with sardines in oil and set at a distance of up to 1 m from the bait. Cameras were set to record still images. Browning cameras were set to ‘RPF-4Shot’ (to take four photos spaced three seconds apart) and the Bushnell cameras were set to take three photos per trigger. Cameras were active for 10 nights and were revisited after five days to replenish the bait and replace the SD cards and batteries in the cameras. All images were screened and species recorded were identified by a combination of volunteers and the authors. Species were recorded to species level, with the exception of small rodents *Rodentia sp*., martens *Martes sp*., and deer *Cervidae sp*. which were classified to group level only, due to the difficulties in correctly identifying these species from camera trap images. All images identified as mink were checked by the authors to confirm the species (European or American).

### Hair tubes

Hair tubes were deployed in pairs approximately 50-100 m apart at each site, giving a total of 10 hair tubes per UTM square. Hair tubes comprised PVC tubes measuring 250 mm length and 75 mm diameter. Two adhesive patches (“mouse glue”; product code: STV182, STV International Ltd., Forge House, Little Cressington, Thetford, Norfolk, IP25 6ND, UK) were attached to the inside of the tube to collect hair from any animal that entered. Hair tubes were fixed vertically, approximately 15 cm from the ground, to either vegetation, such as shrubs or small trees adjacent to the watercourse, or to a metal or wooden rod or stake, with the open end of the tube directed towards the ground. A chicken wing was fixed inside as bait.

Hair tubes were active for 10 nights and were checked daily. When a hair sample was found in the tube, it was removed, stored in ethanol and frozen. The adhesive patches were then replaced and the bait was replenished every 5 days. Hairs were examined under a microscope and identified to species level using characteristics as described in Teerink (1991) and González-Esteban *et al*. (2006).

### Live trapping

Metal cage traps were deployed in pairs approximately 50-100 m apart at each site. Traps were located within a few metres of the hair tube locations, but the trapping was carried out after the hair tube surveys were completed. Traps were baited with sardines in oil and a whole raw hen’s egg. The traps were set for 10 nights, although some traps were closed before the end of the 10 nights, when the water level increased, or when the same individual mink or non-target species was caught repeatedly. Traps were checked daily early in the morning.

When a European mink was captured, it was sedated by a veterinarian, using a mixture of ketamine hydrochloride (Imalgene 1000) and medetomidine (Domitor) administered intramuscularly, and fitted with a subcutaneous microchip for future identification. While the animal was sedated, its sex was determined, various morphometric measurements recorded, and samples taken for genetic analysis - a rooted hair and a piece of cartilage - as part of a larger study to assess the status of the European mink population in the area. Sedation was reversed using atipamoleum (Antisedans) administered intramuscularly in doses equal to the volume of Domitor given. Mink were released at the site of capture within two hours of processing. Other animals captured were released immediately without handling.

### eDNA sampling

Water samples were collected using sterile 500 ml water bottles from the bankside at a reachable distance with complete or near-complete submersion of the bottle beneath the surface. Five 500 ml water replicates (Broadhurst *et al*. 2021) were collected from each site (see description of camera trapping above), within 5-10 m of each other, in close proximity to the camera traps. The first eDNA sampling session was conducted between October 14^th^ – 18^th^ 2019 and then repeated at the same sites between October 24th – 28^th^ 2019, covering the period when the camera traps were active. Samples were filtered on the same day as collection (along with field controls consisting of 500 ml of distilled water to test for cross-contamination during sampling) using sterilised single-use syringes and 0.45 μm Sterivex filters (Merck Millipore, Darmstadt, Germany). The filters containing eDNA were transported in cool boxes with ice packs and then stored at −20°C until DNA extraction.

Four additional 500 ml replicates were taken from a pond within a captive European mink enclosure to test the ability of the method to detect European mink where their presence was known. These samples were transported separately from the field samples.

### eDNA laboratory methods and bioinformatics

DNA was extracted from the filters in a dedicated eDNA clean room following the Mu-DNA protocol (Sellers *et al*. 2018). Field controls were extracted first, followed by the field eDNA samples. After the field samples were processed, the samples from the captive pond were extracted. Eight DNA extraction negative controls (one for each day of extractions) containing only extraction buffers were also included. All surfaces were sterilised with 10% bleach and then washed with 70% ethanol. Small tools were placed in a UV Stratalinker® before, in-between and after extracting each sample to reduce the risk of cross-contamination.

DNA extracts were stored at −20°C until PCR amplification. Eluted eDNA was amplified using the MiMammal 12S primer set (MiMammal-U-F, 5′-GGGTTGGTAAATTTCGTGCCAGC-3′; MiMammal-U-R, 5′-CATAGTGGGGTATCTAATCCCAGTTTG-3′; Ushio *et al*. 2017) targeting a ∼170bp amplicon from a variable region of the 12S rRNA mitochondrial gene with sample-specific multiplex identifier (MIDs) tags. PCR amplification protocols followed Sales *et al*. (2020a) with PCR positive controls (i.e. DNA extraction from a non-target species that is not locally present, the northern muriqui *Brachyteles hypoxanthus* from Brazil included; Broadhurst *et al*. 2021). Sequencing was performed over two runs using the Illumina MiSeq v2 Reagent Kits for 2×150 bp paired-end reads (Illumina, San Diego, CA, USA). The bioinformatic analysis was conducted using OBITools metabarcoding package (Boyer *et al*. 2016) following the protocol described in Sales *et al*. (2020a) and Broadhurst *et al*. (2021). See the Supplementary Material for full details of the laboratory and bioinformatic methods.

### Statistical analysis

We used a Bayesian statistical analysis approach to estimate the detection probabilities of European mink for each method. We estimated the probability of detecting European mink when present throughout the study area using the R (v. 4.1.2; R Core Team, 2021) package ubms (Kellner *et al*. 2021) and STAN software (Carpenter *et al*. 2017) implemented in R Studio (v. 1.2.5042; R Studio Team 2020) with 100,000 iterations, 5 chains and the default burn-in setting of half the number of iterations. The probability of detection of European mink eDNA in a sample replicate was also calculated using the R package eDNAoccupancy (Dorazio & Erickson 2018). Finally, we calculated the probability of detection for the combined use of all possible pairs of methods to assess which combination of methods offered the highest probability of detection.

McNemar’s test was used to compare the efficacy of the two camera models, using the camera model as treatment (Browning) and control (Bushnell).

### Economic cost and time analysis

Costs were calculated for each method, inclusive of all aspects of fieldwork and data collection (purchasing of equipment and consumables for fieldwork, and filtering, shipping and lab costs for eDNA sampling). The time input for each method was estimated, including fieldwork, sample collection, lab work and data analysis. Costs were calculated in Euros (€) and where costs were incurred in GDP (£), these were converted to Euros (€) in October 2021 using the rate of £1 = €1.18, to allow for consistent comparison.

## RESULTS

### Camera trapping

In total, 81,655 camera trap images were recorded and at least 14 mammalian species or groups were detected; red fox *Vulpes vulpes*, European and American mink, Eurasian otter *Lutra lutra*, European pine marten, stone marten *Martes foina*, least weasel *Mustela nivalis*, common genet *Genetta genetta*, domestic cat *Felis catus*, Eurasian beaver *Castor fiber*, red squirrel *Sciurus vulgaris*, small rodent *Rodentia sp*. (classified to group level only), deer *Cervidae sp*., and wild boar *Sus scrofa*. Of all of the images recorded, 938 (1%) images were of mink species. Of these, 85% (n=797) of images were confidently identified as European mink, at 48% (n=12) of sampling sites, in three UTM squares (Fig. 3). Ten images were identified as American mink from one instance at one site. For the remaining 131 images, confident identification of the species was not possible, as the distinguishing features on the face of the mink were not fully visible. The probability of detection of European mink per sampling site by camera trapping was 0.85, with a Bayesian Credibility Interval ranging from 0.51 to 0.99 (Fig. 4b). The average number of days to first detection was 5 (Table S1, Fig. 4a). There was heterogeneity in the number of detections of European mink at a camera trap site, ranging from 1 to 8 (mean=2.8). The activity of European mink at the camera traps was cathemeral, with 49% of images recorded during the night and 51% recorded during daylight hours.

**Fig. 3.**
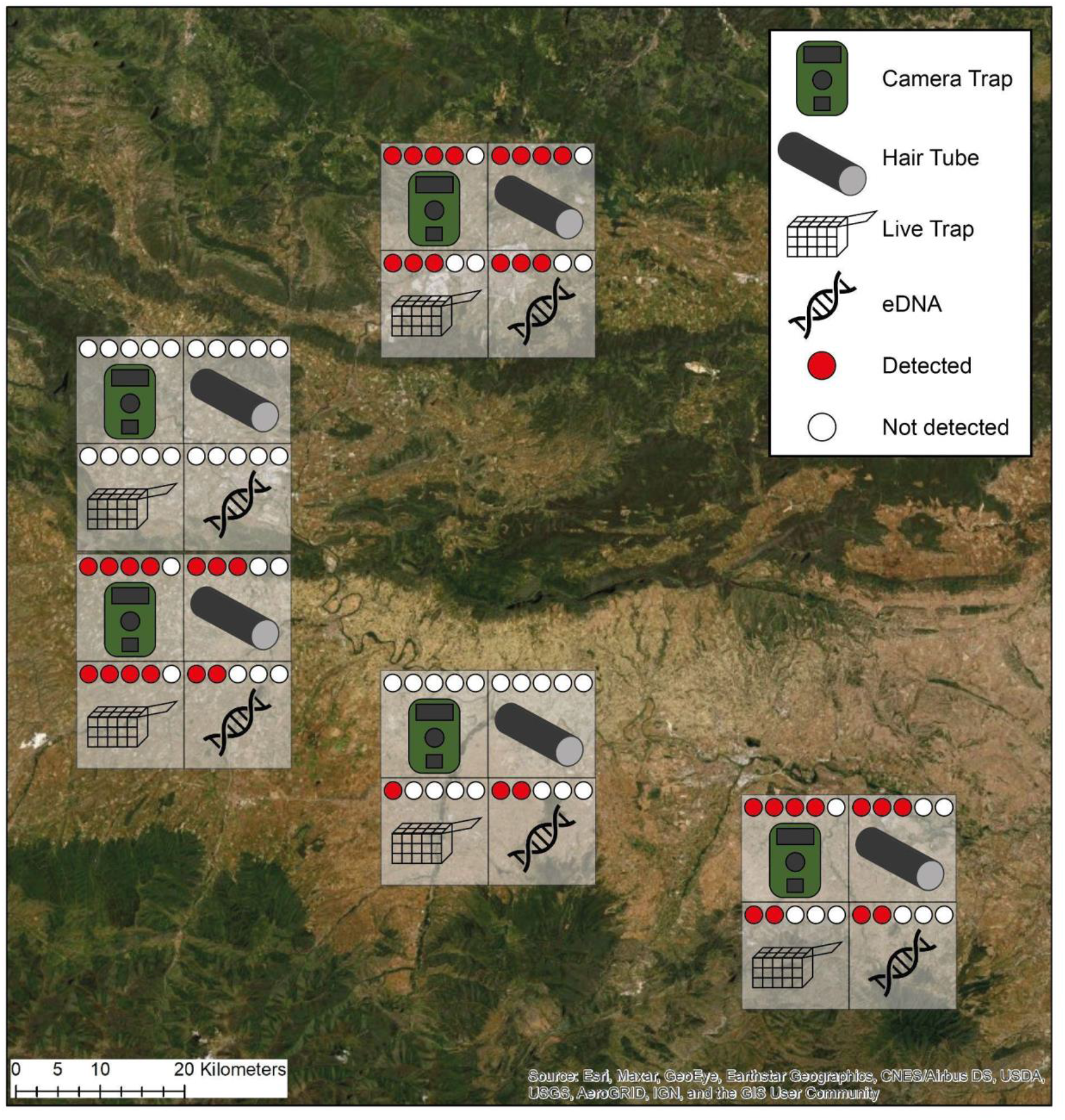
Comparison of the sites in north-eastern Spain where European mink were detected by four surveying methods. The circles by each method represent the individual sampling sites (five per UTM square) where the surveying methods were deployed, and the number of sites where mink were detected or not

**Fig. 4.**
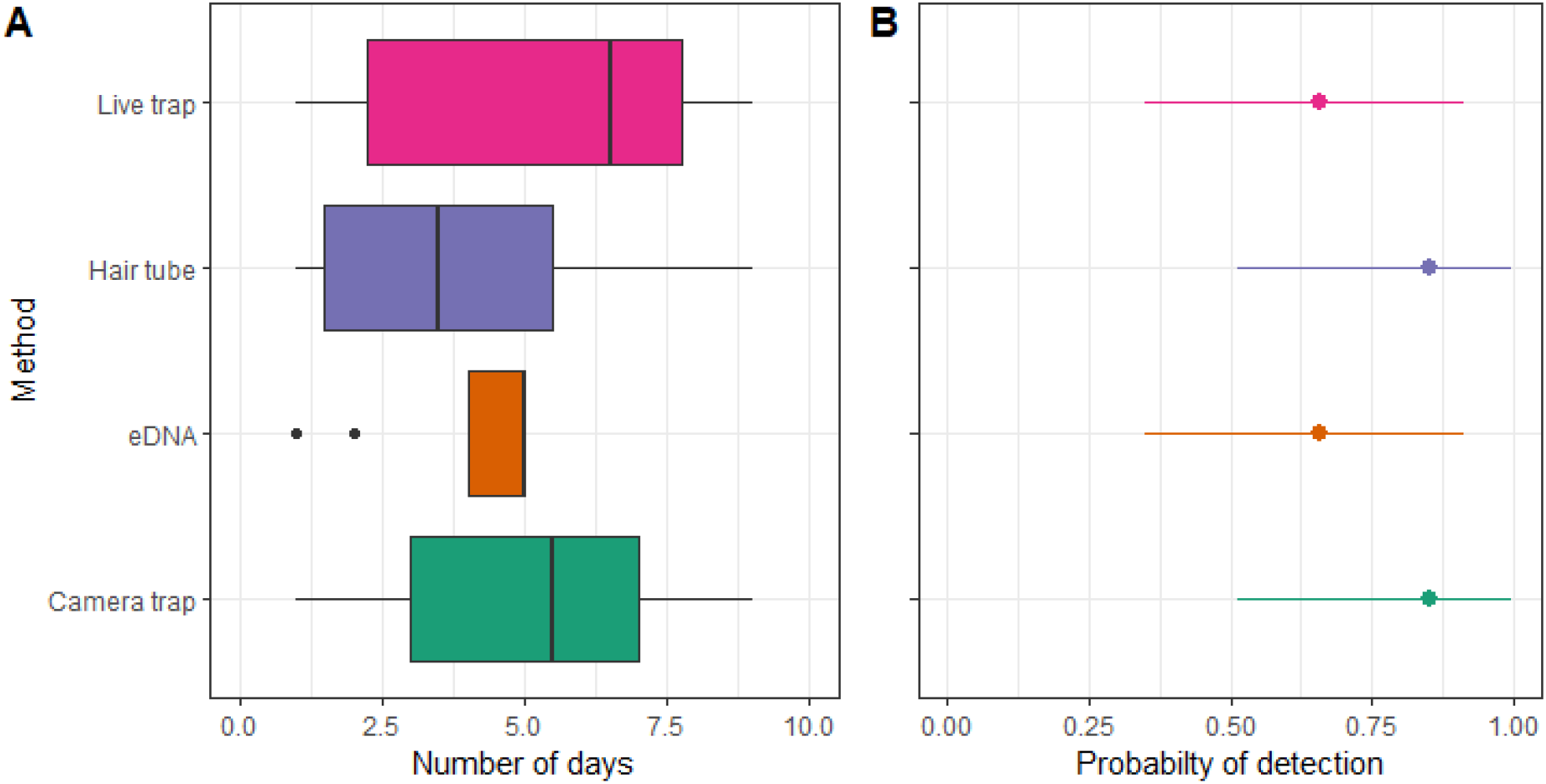
**a** The number of days to the first detection of European mink by four monitoring methods. **4. b** The probability of detection of European mink when present for each method with their 95% Bayesian Credibility Interval. For eDNA sampling, this is represented as the average number of replicates per site (out of a total of 5 replicates per site) to the first detection.

The Browning cameras detected European mink at 12 sites (every site where mink were detected by cameras) and the Bushnell cameras detected European mink at eight sites. Therefore, there were four sites where the Brownings detected European mink and the Bushnells did not, and no sites where the situation was reversed. Nevertheless, this effect was not statistically significant (McNemar’s, *p=*0.134, *χ*^*2*^=2.25, *df*=1).

### Hair tubes

A total of 31 hair samples were collected from the 25 pairs of hair tubes. Of these, 21 samples (68%) were identified as European mink; these were collected from 13 unique hair tubes at 40% (n=10) of sampling sites, in three UTM squares (Fig. 3). The remaining samples were identified as common genet (26%), domestic cat (3%) and brown rat *Rattus norvegicus* (3%). The probability of detection of European mink by hair tubes was 0.85, with a Bayesian Credibility Interval ranging from 0.51 to 0.99 (Fig. 4b). The average number of days to first detection was 3.9 (Table S1, Fig. 4a). The number of detections of European mink at a hair tube site ranged from 1 to 4 (mean=2.1).

### Live trapping

European mink were caught on 12 occasions at 40% (n=10) of sampling sites, in four UTM squares (Fig. 3). Ten individual mink were caught; five males and five females. All individuals were caught on one occasion, with the exception of one female which was caught on three occasions in three different traps. The probability of detection of European mink by live trapping was 0.66, with a Bayesian Credibility Interval ranging from 0.35 to 0.91 (Fig. 4b). The average number of days to first detection was 5.3 (Table S1, Fig. 4a). The number of detections of European mink at a live trapping site ranged from 1 to 2 (mean=1.2).

### eDNA sampling

The two MiSeq sequencing runs yielded a total of 19,091,872 raw sequence reads. Following the quality control and filtering steps outlined previously, 2,210,392 reads were assigned to 24 wild mammalian species and 376,580 reads (17% of total reads) were attributed to European mink. The wild mammalian species detected comprised red fox, European mink, European badger *Meles meles*, otter, weasel, genet, beaver, yellow-necked mouse *Apodemus flavicollis*, brown rat, house mouse *Mus musculus*, edible dormouse *Glis glis*, southern water vole *Arvicola sapidus*, bank vole *Myodes glareolus*, Algerian mouse *Mus spretus*, long-tailed wood mouse *Apodemus sylvaticus*, Eurasian harvest mouse *Micromys minutus*, greater white-toothed shrew *Crocidura russula*, common shrew *Sorex araneus*, aquatic mole *Talpa aquitania*, European rabbit *Oryctolagus cuniculus*, Granada hare *Lepus granatensis*, red deer *Cervus elaphus*, roe deer *Capreolus capreolus* and fallow deer *Dama dama*. All four replicates from the captive pond were positive for European mink. The species was detected at 36% (n=9) of sampling sites across the two sampling sessions, in four UTM squares (Fig. 3). Most mink eDNA detections occurred in the second sampling session (70% in the second sampling session; 30% in the first sampling session; Fig. 5).

**Fig. 5.**
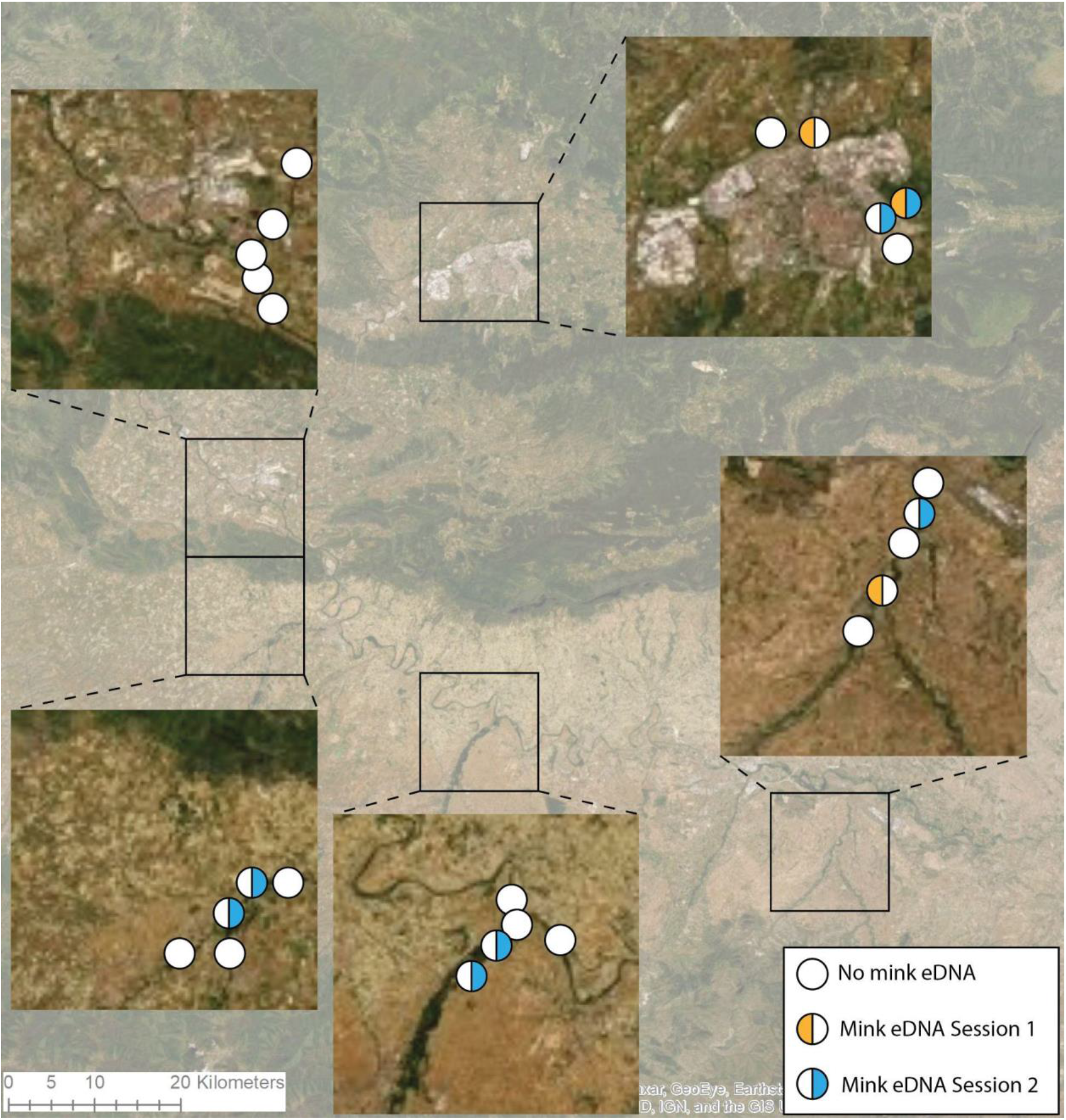
European mink detections by environmental DNA per sampling session (orange representing the first sampling session (October 14-18, 2018) and blue the second (October 24-28, 2018)). Each circle represents the individual sampling sites.

The probability of detection of European mink by eDNA was 0.66, with a Bayesian Credibility Interval ranging from 0.35 to 0.91 when using the R package ubms (Kellner *et al*. 2021) (Fig. 4b), while the probability of detection in a replicate was 0.10 (BCI: 0.04-0.24). The average number of water sample replicates per site to first detection of mink was four (Table S1, Fig. 4b and 4a). All positive sites had an eDNA detection in just one sampling session, with the exception of one site at which mink were detected in both sampling sessions. Out of a total of five water sample replicates taken per site, the number of replicates that were positive for European mink eDNA ranged from 1 to 3, with the majority of sites having one positive replicate (mean=1.3).

### Combined use of two methods

We found that the combination of any two methods offered a very similar probability of detection for European mink (0.77; CI:0.46-0.97; Table S2) with the exception of the use of camera trapping and hair tubes which achieved a higher probability of detection per sampling site when present (0.85; CI:0.51-0.99); however, these two methods did not detect European mink in one UTM square where eDNA and live trapping did.

### Comparison of cost and time

The most expensive method in terms of both cost and time incurred was camera trapping (Table 1). This was due to the considerable cost of purchasing 50 camera traps and the extensive time required to review and classify the camera trap images. eDNA sampling was the second most expensive in terms of cost and time, due to the costs of the lab work (extractions, PCR and sequencing) and the time required for sampling, filtering and lab analysis. Hair tubes and live trapping were the most cost-effective of the four methods (Table 1). This was due to the comparatively shorter time required and the low costs of the field equipment, which was particularly the case for hair tubes, making it the most inexpensive method overall.

**Table 1.** Comparison of cost and time incurred for each monitoring method to detect European mink in north-eastern Spain (including the costs of camera traps and live traps, and all aspects of sample processing and analysis in the field and in the lab).

| **METHOD** | **Method costs (equipment, fieldwork, lab costs)** | **Staff hours (fieldwork, lab work, analysis)** |
| --- | --- | --- |
| **Camera trapping** | € 9,390.00 | 529 |
| **eDNA** | € 7,184.00 | 395 |
| **Hair tubes** | € 500.00 | 124 |
| **Live trapping** | € 2,250.00 | 100 |

## DISCUSSION

Across its global range, the population status and fine-scale distribution patterns of European mink are largely unknown. This knowledge gap is due predominantly to an absence of monitoring (e.g. across Russia and in the Danube Delta; Harrington and Maran, *in press*) but even in areas, such as Spain, where there is considerable monitoring effort, difficulties in assessing population status and distribution arise due to inconsistent approaches among regions and a lack of consensus on how to effectively monitor European mink, and which methods are the most effective and accurate. We aimed to address this gap, by comparing the efficacy of different surveying methods to inform recommendations for a standardised survey approach for European mink.

In this study, all four surveying methods successfully detected European mink, but the probability of detection varied by method, with wide and overlapping Bayesian Credibility Intervals. Camera trapping detected European mink at the most sampling sites and had the joint highest probability of detection, with hair tubes. There was heterogeneity in the number of unique detections by camera per site, with mink detected on more than one occasion at the majority of sites. The camera trap set up worked well in capturing images with a clear view of the animal’s face, allowing reliable discrimination between European or American mink; it was also the only method that detected American mink during the study. Detection of American mink is critical in any European mink monitoring strategy because understanding the distribution of American mink, and early warning of their presence, is critical to informing control efforts to protect European mink (e.g. Põdra *et al*. 2013). American mink are regularly caught in live traps elsewhere, and at other times in this study area, and detected by hair tubes (Põdra, *pers. comm*.); whether or not camera traps are truly more efficient at detecting their presence at low densities requires further study. Further, camera trapping is not species-specific and produced a high proportion of detections of non-target species, with images of European mink comprising just over 1% of the total images recorded. As such, reviewing all of the images took a considerable amount of time, although this could potentially be reduced in future studies through the use of a citizen-science-based initiative or emerging machine learning algorithms to facilitate image classification (Norouzzadeh *et al*. 2018, Green *et al*. 2020, Tabak *et al*. 2019). As well as requiring the highest time input, camera trapping was also the most expensive method, although the cost would be reduced considerably if cameras were not purchased solely for the study and were used for repeat surveys.

This is the first study to confirm the potential of eDNA for monitoring European mink. European mink eDNA was detected at the fewest sampling sites and was typically detected in one water sample replicate out of five. As European mink are a semi-aquatic species, they would seem to be an ideal candidate for eDNA-based monitoring. However, other semi-aquatic carnivores such as the Eurasian otter have proven to be challenging to detect using eDNA, either not being detected at all in sampled areas when presence was known (Sales *et al*. 2020a) or requiring the screening of a comparatively large number of samples to be detected (Broadhurst *et al*. 2021). Carnivores generally have a lower detection rate by eDNA than other mammalian species groups (Broadhurst *et al*. 2021, Lyet *et al*. 2021), with their ecologies (more solitary and wide-ranging) being proposed as the primary reason (Sales *et al*. 2020a). Despite these caveats, eDNA (and live trapping) detected mink in more 10km × 10km squares) than did camera trapping or hair tubes, and eDNA was successful in detecting mink at one sampling site where the species was not detected by any other method, suggesting that eDNA (and live trapping) might be more effective than other methods at a wider landscape scale.

Overall, our results add to other studies which have demonstrated the potential of eDNA metabarcoding for monitoring rare or elusive species (Franklin *et al*. 2019, Sales *et al*. 2020b), but it should be combined with other methods to maximise the chances of detecting European mink, particularly where they are at a low density (Sales *et al*. 2020a). The majority of European mink eDNA samples were detected in the second sampling session (Fig. 5), which may be due to heavy rain which occurred between the first and second sampling session. Lyet *et al*. (2021) reported a similar finding of an increased number of mammalian species detected after rainfall using eDNA and this may be due to increased river flow/transportation and run-off from the terrestrial ecosystem into the adjacent aquatic environment. The cost of eDNA sampling was relatively high and is unlikely to change for future monitoring programmes in the short-term, due to the cost of field and lab materials, and the considerable staff time required for sampling, filtering and lab/bioinformatic analyses. However, a metabarcoding (multi-species) approach was used in this study which is more expensive than a species-targeted approach using either qPCR or droplet digital PCR (ddPCR). Currently, there are no available targeted assays for European mink, however a newly designed targeted eDNA approach could be developed (using the more sensitive ddPCR for example; Mauvisseau *et al*. 2019). Not only would this approach be less expensive, but species-targeted approaches also tend to outperform metabarcoding for detecting a focal species (Harper *et al*. 2019, Wood *et al*. 2020). However, the advantage of the multi-species metabarcoding approach is that it gives additional information on the wider mammalian community (24 species detected compared to 14 species/groups detected by camera traps) and the potential to co-detect the American mink (Broadhurst *et al*. 2021; although American mink were not detected by eDNA in this study), so the additional costs might be outweighed by the additional information provided (depending on the broader aims of the survey). For future monitoring programmes, it will also be important to determine to what extent detections are influenced by eDNA transport and the presence of European mink upstream.

Hair tubes and live trapping detected European mink at the same proportion of sampling sites, although hair tubes had a higher probability of detection than live trapping. Both methods were slightly less effective (i.e. detected mink at fewer sites) than camera trapping, but more effective than eDNA, and both had the advantage of being relatively species-specific (unlike camera trapping and eDNA). However, although in this study, hair tubes performed well (and had the shortest time to detection of all four methods), other longer-term studies comparing hair tube and live trapping surveys carried out in parallel over a larger area in the La Rioja region of Spain than the current study, typically find that live trapping outperforms hair tubes in detecting European mink (M. Põdra, *unpublished data*). A further advantage of live trapping is the immediate identification and rapid removal of any American mink caught. This is not possible with any of the other methods, as it requires a significant amount of time (several weeks/months) to identify American or European mink either through lab analysis of eDNA or hair samples or by reviewing camera trap images, by which time it may be challenging to capture any American mink detected. Live trapping is also the only method which enables the assessment of the condition and health of individual mink (often an important goal when taking into account the critical status of the species) and for PIT tags or radio-collars to be fitted for further monitoring. Nevertheless, live trapping requires experienced personnel, to reduce any risk of injury or incidental mortality to animals captured, and potentially the involvement of a veterinarian depending on the objective of the trapping. As a non-invasive method, hair tubes are a more accessible method, easily implemented by fieldworkers with minimal training. Although not carried out in this study, hair samples can also be genotyped to identify individuals and infer population abundance (e.g. Vergara *et al*. 2014, O’Mahony *et al*. 2017), but this has not yet been validated for European mink in terms of the accuracy of abundance estimates obtained. The cost and time incurred for hair tubes and live trapping was relatively low, thus making them more affordable and feasible in future studies, however, if hair samples were to be genotyped, this would significantly increase the cost and time of this method.

The detectability of a species by different methods is influenced by behavioural traits which dictate the individual response of an animal to the method (Merrick & Koprowski 2017). Bold, active, exploratory or aggressive individuals might be more likely to explore and be detected by novel objects, such as traps and hair tubes (Carter *et al*. 2012), whilst these methods may fail to detect less active, neophobic or wary individuals (Merrick & Koprowski 2017). If this were the case for European mink, live trapping and hair tubes may fail to detect neophobic individuals, and these individuals may only be detected by eDNA sampling and camera trapping. In this instance, using a combination of methods may maximise the detection of a range of individuals and behavioural types. Our results also suggest that the relative efficacy of the survey methods tested here may differ according to the abundance of mink present, and may offer different advantages over different scales (for example, eDNA might be particularly useful for detecting mink presence over much larger areas prior to identifying sites for more detailed monitoring; Sales *et al*. 2020a). However, since we were only able to compare methods over a relatively small spatial scale (five discrete UTM squares), further study over a broader scale is required to verify the patterns observed here.

### Recommendations and conclusion

For future surveys or monitoring programmes for European mink, the choice of method(s) should be dictated by the exact objectives of the study and ideally use a combination of at least two methods, in order to maximise detectability of mink.

Our results suggest that both cameras and hair tubes provide effective non-invasive methods, with cameras detecting mink at the highest number of sampling sites and providing additional data on other species, although being considerably more expensive and slower to the first detection of European mink. However, to ensure detection of mink at low population densities, our results also suggest that it is important that these methods are combined with either live trapping or eDNA sampling (depending on the aim of the survey) to accurately map distribution and collect reliable occupancy data. Live trapping is a well-established method for European mink which has been used extensively in Spain, and provides individual-level data that other methods cannot that are crucial for assessing population health and viability. eDNA has potential for use over large landscape scales to determine broad distribution patterns of European mink (and thus to refine current national and globe range estimates), whilst also providing additional presence data on a wide range of mammalian species. For eDNA sampling, we would recommend taking no fewer than five water sample replicates per site and there would likely need to be an increase in spatial and temporal resolution of samples and replicates, and the trialling of a species-specific eDNA assay (e.g. ddPCR) in comparison to multi-species (metabarcoding). For camera trapping, we would recommend using two cameras at each sampling site to maximise the chances of achieving a clear image of the mink’s face, in order to distinguish between European and American mink, and to account for possible malfunctions of the cameras. As the outlook for European mink remains perilous across its range, implementing the surveying methods described here can provide critical information on the distribution and conservation status of extant populations. This will be particularly important to inform crucial conservation interventions and facilitate monitoring of proposed reintroductions.

## Acknowledgements

The live trapping and hair tube surveys were supported by La Rioja Regional Government, the Ministry for the Ecological Transition and the Demographic challenge and Tragsatec. The eDNA component of this work was funded by grants from the Vincent Wildlife Trust, People’s Trust for Endangered Species and a University of Salford Internal Research Award given to ADM. We thank the members of the Molecular Ecology Group in Salford for advice on laboratory work and bioinformatics. We are grateful to all of the volunteers who assisted with classification of camera trap images.

## Declarations

The authors declare no competing interests.

This study involved live trapping European mink, and handling of the animals under anaesthesia – these procedures were carried out as part of a larger on-going TRAGSATEC study of European mink in Spain and were carried out under licence from regional governments.

## Data availability

The datasets generated during the current study are available from the corresponding author upon reasonable request.

## SUPPLEMENTARY INFORMATION

### Laboratory Methods and Bioinformatics

PCRs were conducted in triplicates to reduce bias in individual reactions and the replicates were pooled prior to library preparation. Amplification was validated using 1.2% agarose gel electrophoresis stained with GelRed (Cambridge Bioscience). PCR positive controls (i.e. DNA extraction from a non-target species that is not locally present, the northern muriqui *Brachyteles hypoxanthus* from Brazil, at a concentration of 0.05 ng/µl; Broadhurst et al. 2021) were included. eDNA samples were equally distributed into four libraries and across two sequencing runs, with replicated extraction, positive and captive pond controls in each library. A left-sided size selection was performed using 1.1x Agencourt AMPure XP (Beckman Coulter) and Dual-Index adapters (Illumina) were added to each library using KAPA HyperPrep kit (Roche). Each library was then quantified by qPCR using NEBNext qPCR quantification kit (New England Biolabs) and pooled in equimolar concentrations of 9pM. Libraries were sequenced in two runs using an Illumina MiSeq v2 Reagent Kit for 2×150 bp paired-end reads (Illumina, San Diego, CA, USA).

The bioinformatic analysis was conducted using OBITools metabarcoding package (Boyer *et al*. 2016) following the protocol described in Sales *et al*. (2020a) and Broadhurst *et al*. (2021). Briefly, the quality of the reads were assessed using FASTQC (Andrews 2015), a filter was used to select fragments of 140-190bp and to remove reads with ambiguous bases using obigrep, followed by a sequence clustering using SWARM (Mahé *et al*. 2015) and a taxonomic assignment conducted using ecotag against a custom database (Sales *et al*. 2020a). An additional conservative filtering procedure was conducted to exclude MOTUs/reads originating from putative sequencing errors or contamination in order to avoid false positives. First, to account for the occurrence of tag-jumps between tagged amplicons (Schnell *et al*. 2015), the frequency of tag-jumping was calculated using the positive control (PC, *B. hypoxanthus*) according to Broadhurst *et al*. (2021). This frequency was taken off all MOTUs and the PC was removed. Then, to remove putative contaminants, the maximum number of reads recorded for a MOTU in one of the negative controls (whether this be a field collection blanks, extraction blanks or PCR negative controls) was removed from all samples for each MOTU. Only reads associated with European mink with a best identity of > 0.98 (Sales *et al*. 2020a) were retained for the purposes of this study. These read counts in a water replicate at a sampling site were then converted into binary presence-absence data for downstream analyses (Broadhurst *et al*. 2021).

**Table S1.** Comparison of four surveying methods used for detecting European mink, based on a total of 25 sample sites within 5 UTM squares (5 sites per square). *For eDNA sampling, this is represented as the average number of replicates per site (out of a total of 5 replicates per site) to the first detection.

| **Method** | **Proportion of sampling sites European mink detected** | **Proportion of UTM squares European mink detected** | **Probability of detection (Bayesian Credibility Interval)** | **Average number of days until first detection of European mink** |
| --- | --- | --- | --- | --- |
| **Camera trapping** | 48% (n=12) | 60% (n=3) | 0.85 (0.51-0.99) | 5 |
| **Hair tubes** | 40% (n=10) | 60% (n=3) | 0.85 (0.51-0.99) | 3.9 |
| **Live trapping** | 40% (n=10) | 80% (n=4) | 0.66 (0.35-0.91) | 5.3 |
| **eDNA** | 36% (n=9) | 80% (n=4) | 0.66 (0.35-0.91) | 4* |

**Table S2.**
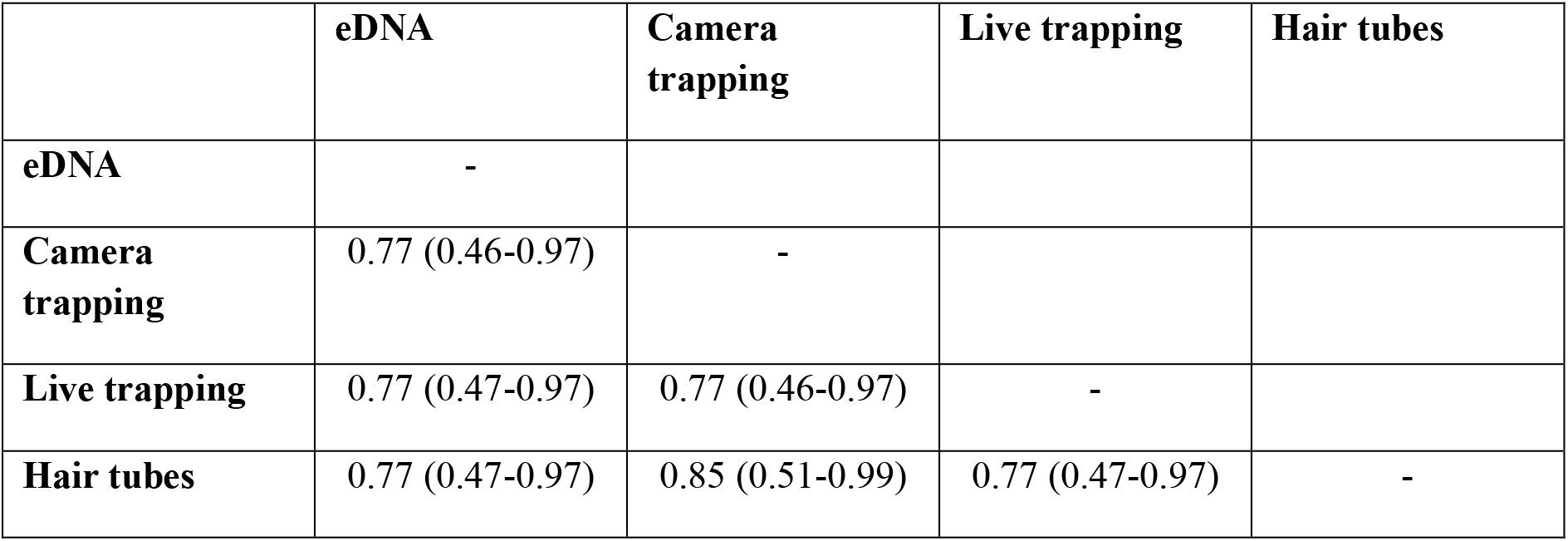
The probability of detection for the combined use of two surveying methods to detect European mink in north-eastern Spain.

|  | **eDNA** | **Camera trapping** | **Live trapping** | **Hair tubes** |
| --- | --- | --- | --- | --- |
| **eDNA** | - |  |  |  |
| **Camera trapping** | 0.77 (0.46-0.97) | - |  |  |
| **Live trapping** | 0.77 (0.47-0.97) | 0.77 (0.46-0.97) | - |  |
| **Hair tubes** | 0.77 (0.47-0.97) | 0.85 (0.51-0.99) | 0.77 (0.47-0.97) | - |

## Notes

### Competing Interest Statement

The authors have declared no competing interest.

### Summary of Updates

I've only edited the author list, as one author was missing

